# TCGAbiolinksGUI: A graphical user interface to analyze GDC cancer molecular and clinical data

**DOI:** 10.1101/147496

**Authors:** Tiago C Silva, Antonio Colaprico, Catharina Olsen, Gianluca Bontempi, Michele Ceccarelli, Benjamin P Berman, Houtan Noushmehr

## Abstract

**Background:** The GDC (Genomic Data Commons) data portal provides users with data from cancer genomics studies. Recently, we developed the R/Bioconductor TCGAbiolinks package, which allows users to search, download and prepare cancer genomics data for integrative data analysis. The use of this package requires users to have advanced knowledge of R thus limiting the number of users.

**Results:** To overcome this obstacle and improve the accessibility of the package by a wider range of users, we developed *TCGAbiolinksGUI* that uses shiny graphical user interface (GUI) available through the R/Bioconductor package.

**Conclusion:** The TCGAbiolinksGUI package is freely available within the Bioconductor project at http://bioconductor.org/packages/TCGAbiolinksGUI/. Links to the GitHub repository, a demo version of the tool, a docker image and PDF/video tutorials are available at http://bit.do/TCGAbiolinksDocs.

## Background

The National Cancer Institute’s (NCI) Genomic Data Commons (GDC), a data sharing platform that promotes precision medicine in oncology, provides a rich resource of molecular and clinical data of almost 13,000 tumor patient samples across 38 different cancer types and subtypes. The platform includes data from The Cancer Genome Atlas (TCGA) and Therapeutically Applicable Research to Generate Effective Treatments (TARGET). The data, which is publicly available, have been utilized by researchers to make novel discoveries and/or validate important findings. To enhance these findings, several important bioinformatics tools to harness genomics cancer data were developed, many of them belonging to the Bioconductor project [1]. Among those tools is our tool TCGAbiolinks [2], which was developed to facilitate the analysis of TCGA data by incorporating the query, download and processing steps within the Bioconductor project [1]. This tool allows users to integrate TCGA data with Bioconductor packages thus harnessing a wealth of statistical methodologies for biologically derived data. In addition, it provides integrative methodologies to perform several important downstream analyses, such as DNA methylation and Gene expression integration. A full detailed comparison between TCGAbiolinks and other bioinformatics tools to analyze TCGA data was previously detailed in our report in which we highlight key advantages of using TCGAbiolinks [2]. Although TCGAbiolinks is a suitable R package for most data analysts with a strong knowledge and familiarity with R specifically those who can comfortably write strings of common R commands, we developed TCGAbiolinksGUI to enable user access to the methodologies offered in TCGAbiolinks and to give users the flexibility of point-and-click style analysis without the need to enter specific arguments. TCGAbiolinksGUI takes in all the important features of TCGAbiolinks and offers a graphics user interface (GUI) thereby eliminating any need to familiarize TCGAbiolinks’ key functions and arguments. Tutorials via online documents and YouTube video instructions will assist end-users in taking full advantage of TCGAbiolinks. Here we present TCGAbiolinksGUI an R/Bioconductor package which uses the R web application framework shiny [3] to provide a GUI to process, query, download, and perform integrative analyses of TCGA data.

## Implementation

### Infrastructure

The TCGAbiolinksGUI user interface was created using Shiny, a Web Application Framework for R, and uses several packages to provide advanced features that can enhance Shiny apps, such as shinyjs to add JavaScript actions [4], shinydashboard to add dashboards [5] and shinyFiles [6] to provide access to the server file system. The following R/Bioconductor packages are used as back-ends for the data retrieval and analysis: TCGAbiolinks [2] which allows to search, download and prepare data from the NCI’s Genomic Data Commons (GDC) data portal into an R object and perform several downstream analysis; ELMER (Enhancer Linking by Methylation/Expression Relationship) [7, 8] which identifies DNA methylation changes in distal regulatory regions and correlate these signatures with the expression of nearby genes to identify transcriptional targets associated with cancer; ComplexHeatmap to visualize data as oncoprint and heatmaps, pathview [10] which offers pathway based data integration and visualization; and maftools [11] to analyze, visualize and summarize MAF (Mutation Annotation Format) files.

### Graphical user interface design

The user interface has been divided into three main GUI menus. The first menu defines the acquisition of GDC data. The second defines the analysis steps which subdivides according to the molecular data types. And the third is dedicated to harnessing integrative analyses. We present below a brief description of each menu and their features that can be accessed through a side panel (see figure 1):

**Figure 1.**
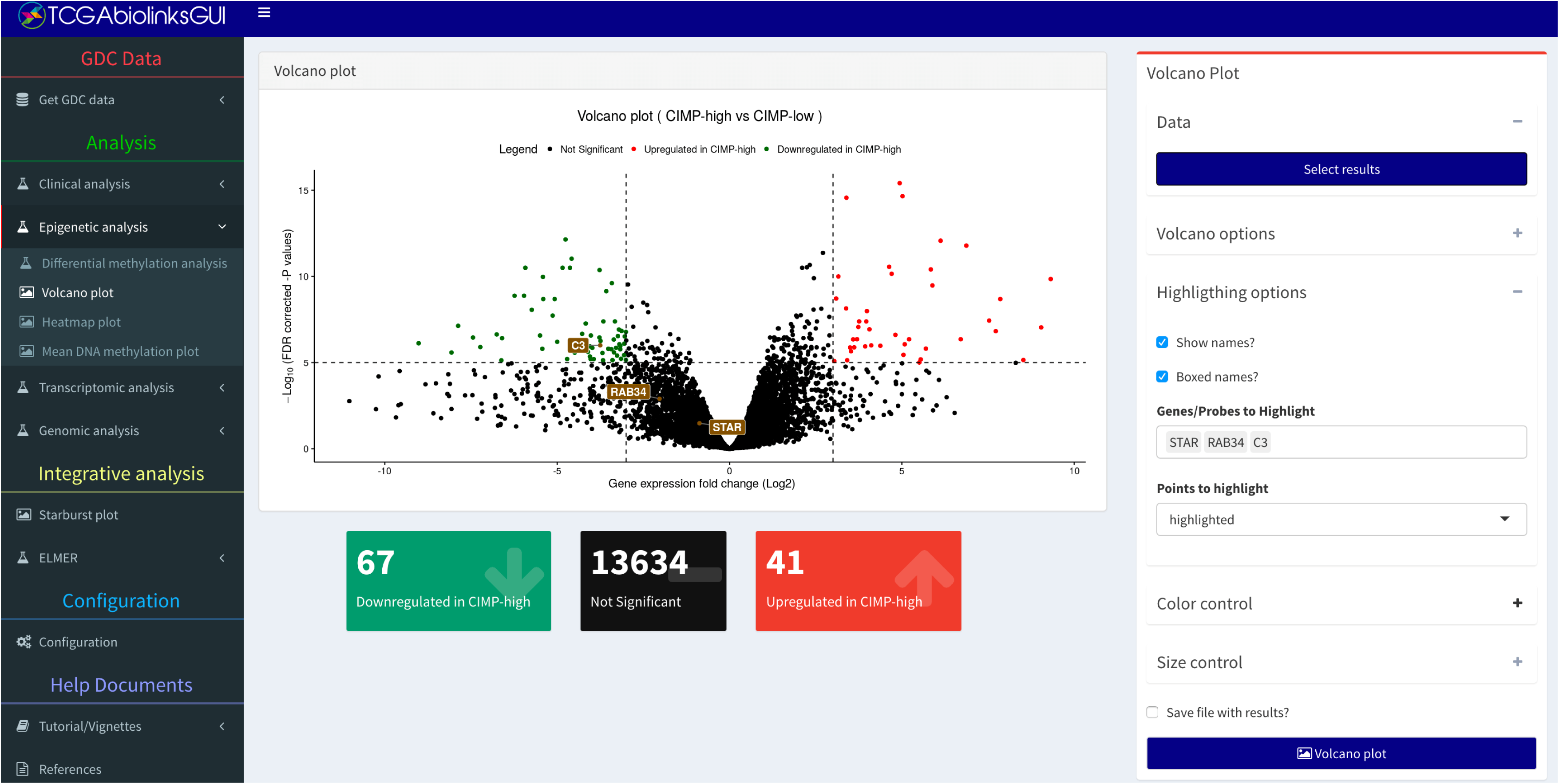
The volcano plot menu of TCGAbiolinksGUI. The panel on the left shows the menus divided by different analyses, the panel on the right shows the controls available for the menu selected. In the center is a volcano plot window from the analysis menu. It is possible to control the colors, to change cut-offs and to export results into a CSV document.

- **GDC Data:** Provides a guided approach to search for published molecular subtype information, clinical and molecular data. In addition, it downloads and processes the molecular data into an R object that can be used for further analysis.
- **Clinical analysis:** Performs survival analysis to quantify and test survival differences between two or more groups of patients and draws survival curves with the ‘number at risk’ table, the cumulative number of events table and the cumulative number of censored subjects table using the R/CRAN package survminer [12].
- **Epigenetic analysis:** Performs a Differentially methylated regions (DMR) analysis, visualizes the results through both volcano and heatmap plots, and visualizes the mean DNA methylation level by groups.
- **Transcriptomic analysis:** Performs a Differential Expression Analysis (DEA), and visualizes the results through both volcano and heatmap plots. For the genes found as upregulated or downregulated an enrichment analysis can be performed and pathway data can be integrated [10].
- **Genomic analysis:** Visualize and summarize the mutations from MAF (Mutation Annotation Format) files through summary plots and oncoplots using the R/Bioconductor maftools package [9, 11].
- **Integrative analysis:** Integrate the DMR and DEA results through a starburst plot. Also, using the DNA methylation data and the gene expression data the R/Bioconductor ELMER package can be used to discover functionally relevant genomic regions associated with cancer [7, 8].

### Documentation

We provide a guided tutorial for users via a vignette document which details each step and menu function available at http://bit.do/TCGAbiolinksDocs, via online documents available at http://bit.ly/TCGAbiolinks_PDFTutorials, and via YouTube video instructions showing step by step how each menu works available at http://bit.ly/TCGAbiolinksGUI_videoTutorials, which assist end-users in taking full advantage of TCGAbiolinksGUI. A demonstration version of the tool is available at http://tcgabiolinks.fmrp.usp.br:3838/. Users are encouraged to report and file bug reports or feature requests via our GitHub repository BioinformaticsFMRP/TCGAbiolinksGUI/issues.

### Docker container

To further simplify the usability and accessibility of our tool, we provide a docker image compatible with most popular operating system available at https://hub.docker.com/r/tiagochst/tcgabiolinksgui/. This file allows users to run TCGAbiolinksGUI without the need to install associated dependencies or configure system files, common steps required to run R installations and load R/Bioconductor packages.

## Results and Discussion

To provide the users with insights into the usability of our TCGAbiolinksGUI, 1) we compare with other bioinformatics tools currently published in the field; 2) we provide a use-case that allows users a step-by-step guide to analyzing their own cancer molecular data.

### Comparison of alternative software

Web tools used for cancer data analysis might be classified into two broad groups. The first group only provides an interface to existing software analysis tools. The Galaxy project (https://galaxyproject.org/), which is an open, web-based platform for accessible, reproducible, and transparent computational biomedical research, is an example of such a tool that belongs to this group. The other group is composed of exploratory tools mainly focused on the visualization of processed data and pre-computed results. The cBioPortal project [13, 14], by providing several visualizations for mining the TCGA data, is an example of a tool that falls within this classification.

If one were to classify TCGAbiolinksGUI, it would belong to the first group. Compared to the Galaxy project, TCGAbiolinksGUI offers an open platform which improves the accessibility of R/Bioconductor packages, allowing users an advantage to integrate their features with existing bioconductor packages without the need to go beyond the R/Shiny frameworks as a common feature from the Galaxy project, which requires the interface elements to be structured through XML files [15]. In addition, going beyond the R/Bioconductor environment requires more software dependencies which make the process to install Galaxy to use R/Bioconductor packages laborious. On the other hand, compared to cBioPortal, TCGAbiolinksGUI allows users to perform deep integrative analysis by comparing different subtypes of data (i.e. performing an integrative analysis to compare breast cancer samples with a mutation on FOXA1 gene compared to wild-type samples using DNA methylation, gene expression, and motif enrichment analysis on genomic regions of interest). Although cBioPortal offers these features, it would require users to process each step independently and download outside of cBioPortal in order to perform such integrative analysis.

### Use case

As an example of integrative analysis available through TCGAbiolinksGUI, we will download Lung Squamous Cell Carcinoma (LUSC) gene expression and DNA methylation data from Genomic Data Commons (GDC) data portal [16] which will be used to perform the R/Bioconductor ELMER analysis. This package allows one to identify DNA methylation changes in distal regulatory regions and correlate these signatures with expression of nearby genes to identify transcriptional targets associated with cancer. For these distal probes correlated with a gene, a transcription factor motif analysis is performed followed by expression analysis of transcription factors to infer upstream regulators. However, instead of using all LUSC samples available in GDC, which would take hours to download and execute the analysis, the steps will be done with a subset of the data. To be able to execute the workflow in a feasible time, the first step requires the download of two samples, while for the analysis steps we provide a link to the processed data of more than 200 samples and only within chromosome 1. A more detailed version of this use case is available in https://bioinformaticsfmrp.github.io/Bioc2017.TCGAbiolinks.ELMER/index.html. The download of molecular data is accessible through the GDC data menu. To help users to use the correct fields, commonly available in the GDC data portal, the required or optional fields will be shown while the unnecessary ones will be hidden. For example, for any search, it is required to specify at least the GDC project, the data category and the data type. For the gene expression data, the field work-flow type is required, while for the DNA methylation data, the field platform is required. Also, users have the option to filter samples by sample type (Primary Solid tumor, Solid tissue normal), barcode or based on Clinical data (i.e only female samples, long or short-term survivors, etc.).

After the data is searched, it can be downloaded and prepared into a SummarizedExperiment object [17], which will hold data and metadata in an organized structure. Figure 2 highlights the field values for the downloaded gene expression data for two primary solid tumor samples (”TCGA-34-5231-01”,”TCGA-77-7138-01”) while figure 3 shows the fields for the downloaded DNA methylation data.

**Figure 2.**
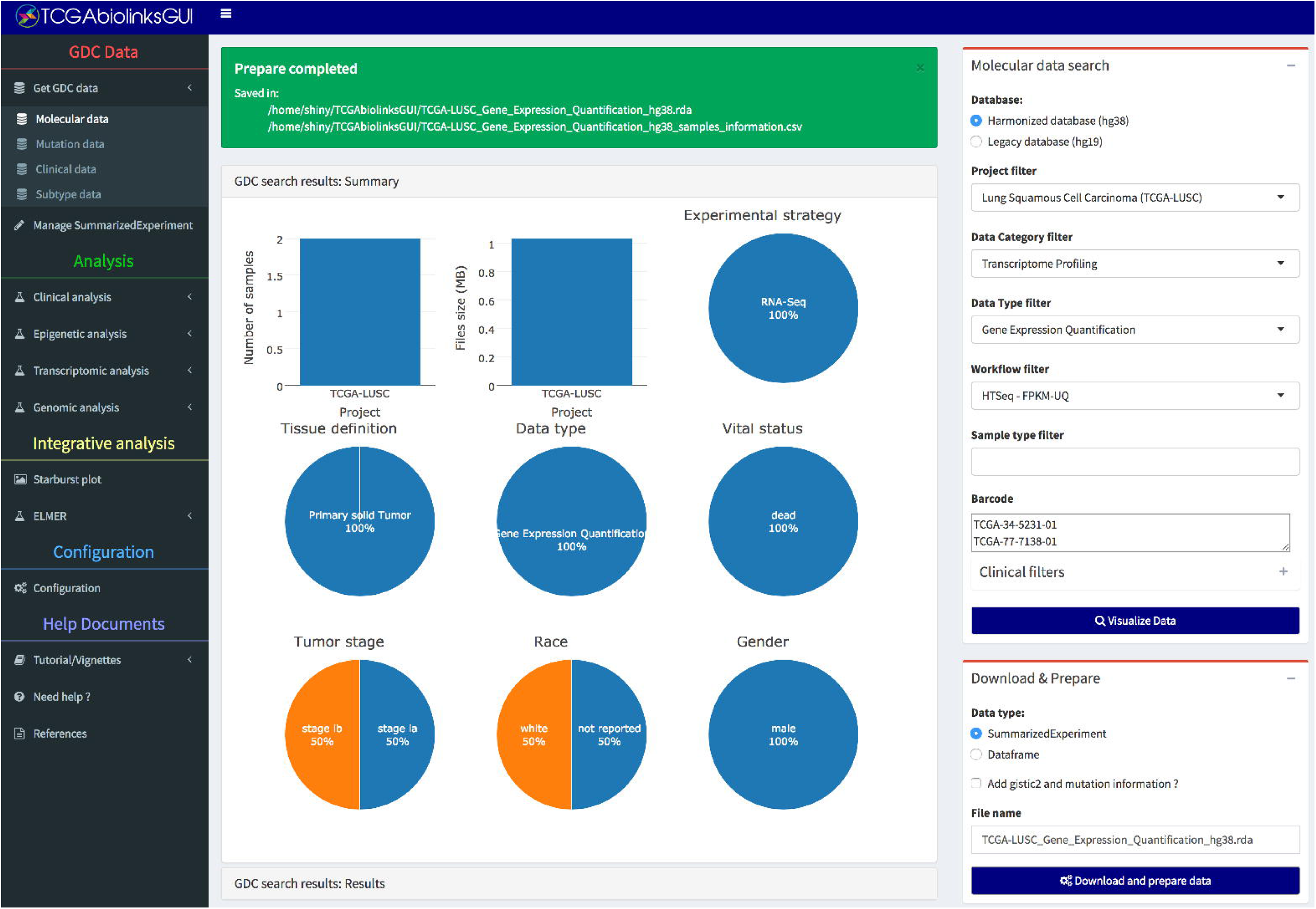
Molecular data download: Gene expression. Search, download and prepare into an R object of gene expression data for two TCGA-LUSC samples (”TCGA-34-5231-01”,”TCGA-77-7138-01”). The Summarized Experiment object created is saved as an RData file (TCGA-LUSC-Gene Expression Quantification hg38.rda).

**Figure 3.**
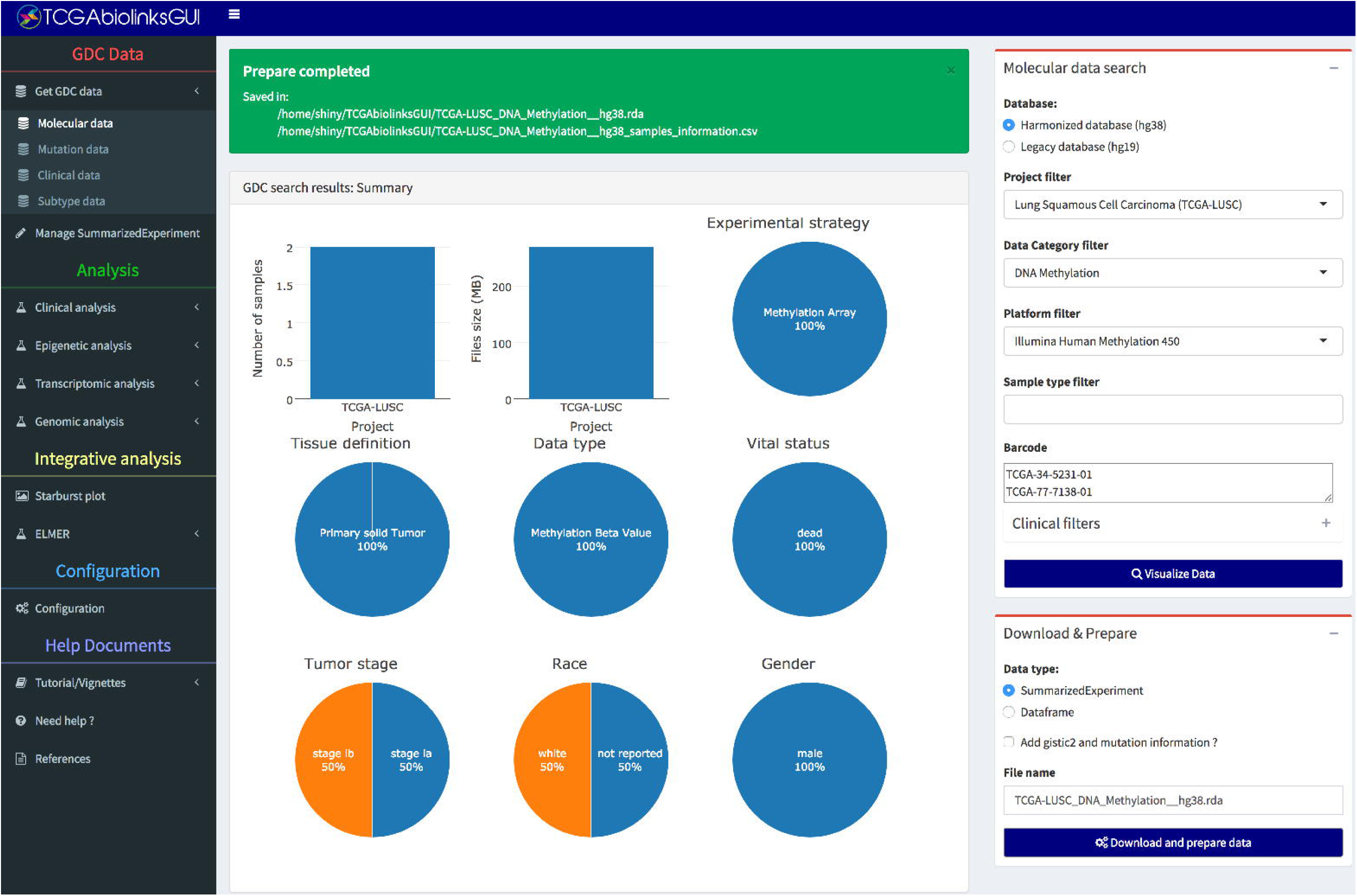
Molecular data download: DNA methylation. Search, download and prepare into an R object of DNA methylation for two TCGA-LUSC samples (”TCGA-34-5231-01”,”TCGA-77-7138-01”). The Summarized Experiment object created is saved as an RData file (TCGA-LUSC-DNA methylation hg39.rda).

For the ELMER analysis step, as explained previously, we will use a pre-processed DNA methylation object containing 268 samples available at http://bit.ly/LUSC_DNAmet and a pre-processed gene expression object containing 234 samples available at http://bit.ly/LUSC_exp. These objects will be used to create an ELMER input data, a MultiAssayExperiment (MAE) [18] object which is a single object able to handle data with different features (i.e. DNA methylation with probes as features and Gene expression data with genes as features). ELMER verifies if a sample has both DNA methylation and gene expression, otherwise, it is removed from the final object, and it keeps the distal probes (those at least *±*2*Kb* pair of the transcription start sites (TSSs)). Upon a successful creation, a message with the object path will be shown (Figure 4).

**Figure 4.**
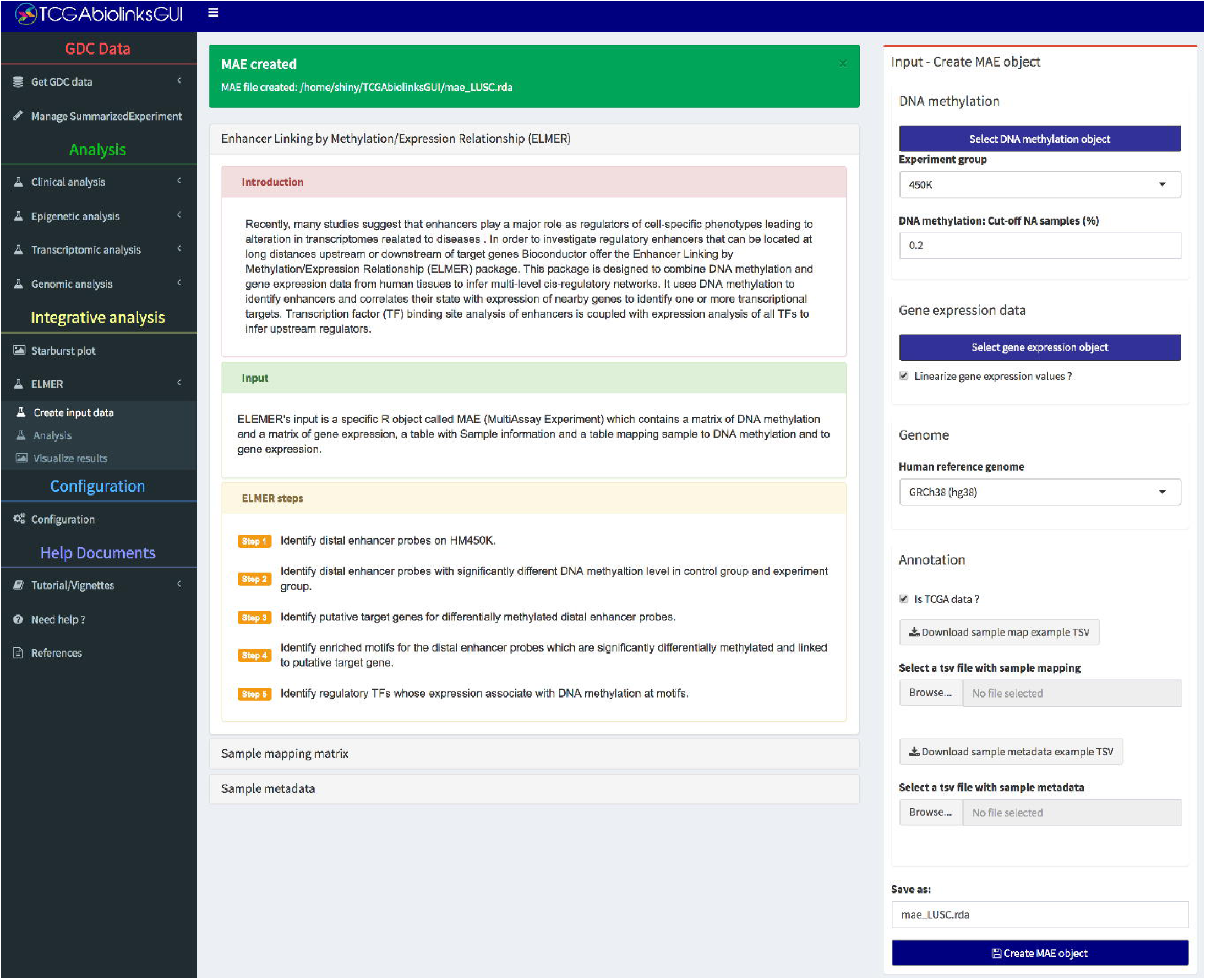
ELMER input data menu. Creating a MAE (MultiAssayExperiment) object only with samples that have both DNA methylation and gene expression data. Probes will be filtered to remove those having NA (empty values) for more than 20% of the samples and only those in distal regions (*±*2*Kb* away from TSS) will be kept.

For the ELMER analysis (Figure 5), the first input required is an MAE object, followed by the selection of the groups to be compared based on the metadata available in the object; in our example, we will compare “Primary Solid Tumor” samples versus “Normal tissue Samples”. For the first step which identifies significant differently methylated probes, we will set to find “Probes hypomethylated in Primary Solid Tumor compared to Solid Tissue Normal”. For the second step (the most time consuming), which identifies anti-correlated gene expression and DNA methylation levels and corrects the significance using a permutation approach, we will change the value of the field “Number of permutations” to 100, “Raw P-value cut-off” to 0.05 and “Empirical P value cut-off” (corrected p-value using the permutation approach) to 0.01.

**Figure 5.**
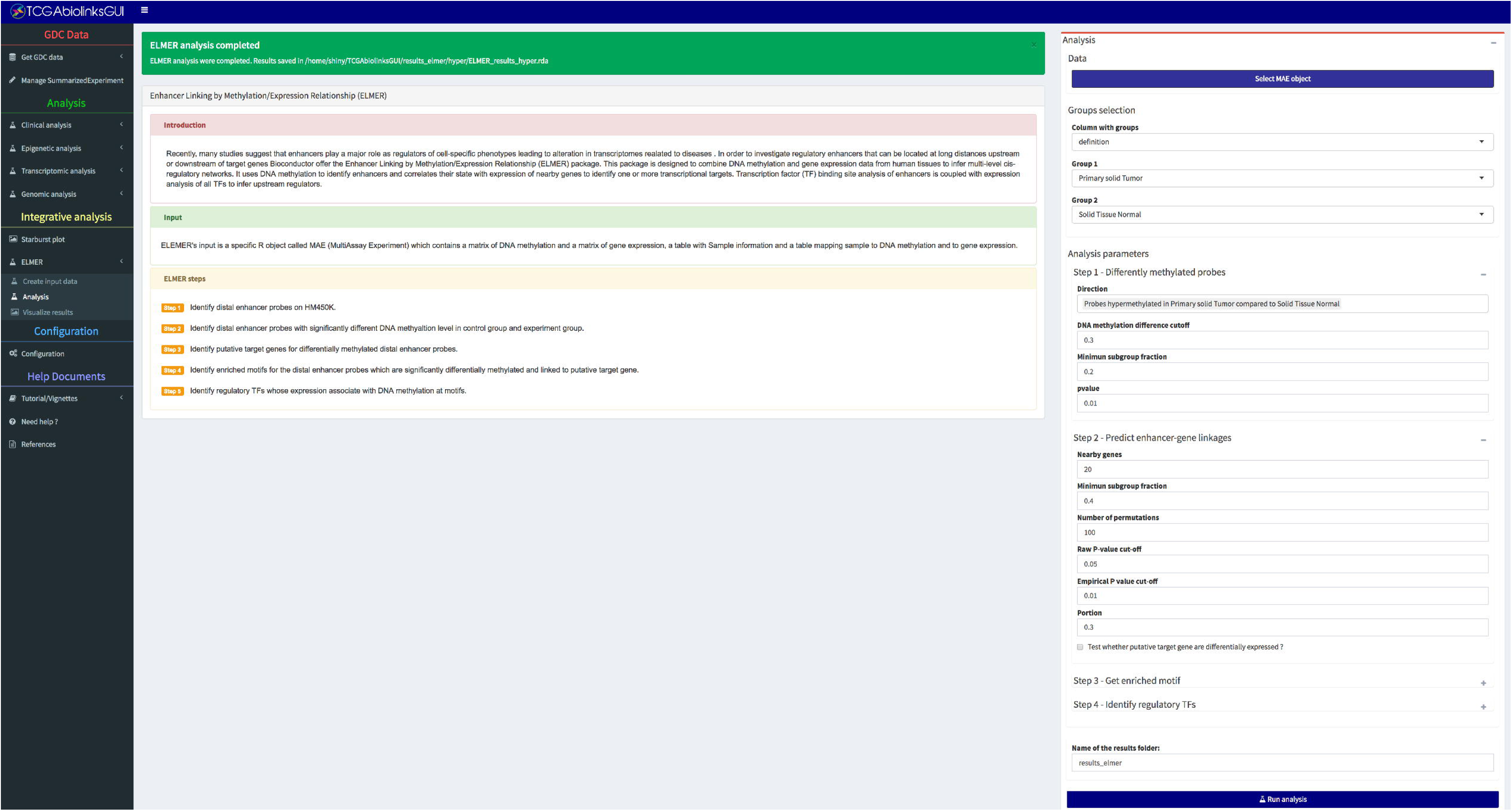
ELMER analysis menu. Result of ELMER analysis for Primary Solid Tumor vs Solid Tissue Normal of Lung Squamous Cell Carcinoma (LUSC) samples. ELMER searches for probes hypomethylated in Primary Solid Tumor and correlate those regions with the expression of their 20 nearest genes. A motif enrichment analysis is performed on the pairs anti-correlated (loss of DNA methylation and gain of expression in a distal gene), followed by the inference of candidate regulatory transcription factors. If any enriched motifs are found, an object with the results is saved to be later visualized through plots and tables.

If there are no results found, ELMER will show an error message. However, if it was able to find candidate regulatory transcription factors that are associated with hypomethylated regions with the potential to regulate distal genes, ELMER will save an object to visualize the results as either tables or plots (Figure 6)

**Figure 6.**
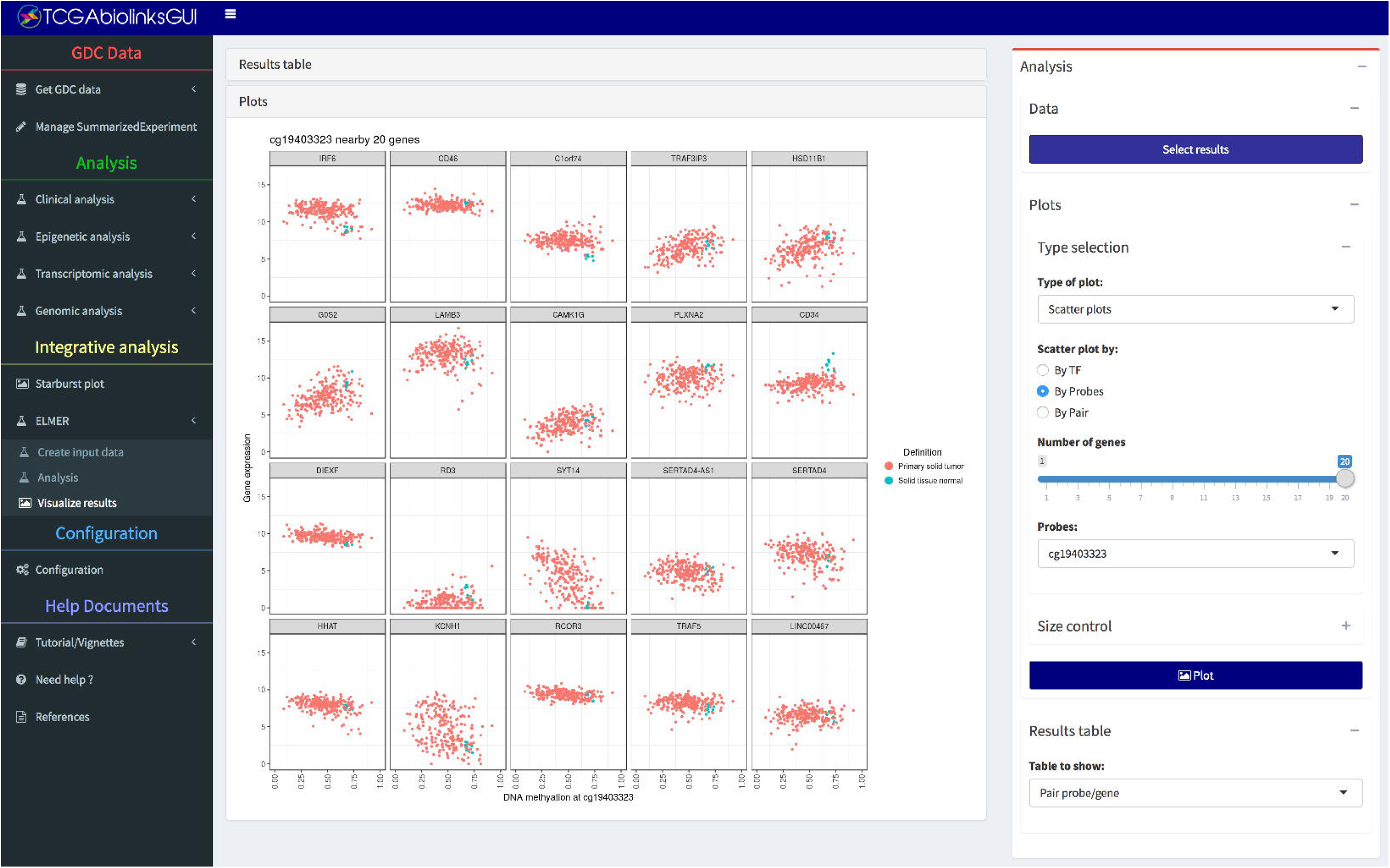
ELMER visualize results menu. With the result object of ELMER analysis, users are able to visualize scatter plots (expression of 20 nearest genes of a probe vs its DNA methylation level, TF expression vs mean DNA methylation of probes with a given motif, gene expression vs probe DNA methylation level), schematic plots (visualization of pairs of probe and genes), odds ratio (OR) plot (visualization of the motif enrichment analysis results), TF ranking plot (ranking of significance for all TF expression vs mean DNA methylation on the paired probes (binding sites)). Through tables, the user is able to visualize the list of significant differentially methylated probes, significant anti-correlated pairs of genes and probes, enriched motifs and candidate regulatory TFs.

## Conclusion

TCGAbiolinksGUI was developed to provide a user-friendly interface of our popular TCGAbiolinks package. TCGAbiolinksGUI is designed specifically for 1) the least experienced R user, who will be able to import GDC data and perform R/Bioconductor analysis without the need to know how to program in R; 2) the most experienced R user, who will be able to execute several of the R/Bioconductor functions by not needing to write several lines of code. For the R/Bioconductor developers, the package has an extensible design in allow developers to add new features by modifying a few lines of the main code and adding a file with user interface elements on the client side and a file with their control on the server side. Also, TCGAbiolinksGUI supports the most updated R/Bioconductor data structures (i.e. SummarizedExperiment and MultiAssayExperiment) which allows to handle data and metadata into one single object and validates several integrity requirements. Thereby, TCGAbiolinksGUI package allows data handling to be as efficient as possible and thereby limits and avoids user errors in data manipulation such as sample removal that involves also metadata deletion.

Finally, several efforts to understand genomic and epigenomic alterations associated with tumor development has been made in the last few years, which presents several bioinformatics challenges, such as data retrieval and integration with clinical data and other molecular data types. This package will allow end-users to facilitate the mining of cancer data deposited in GDC, in hopes to aid in analyzing and discovering new functional genomic elements and potential therapeutic targets for cancer.

## Availability and requirements

Project name: TCGAbiolinksGUI
Project home page: http://bioconductor.org/packages/TCGAbiolinksGUI/
Project GitHub: https://github.com/BioinformaticsFMRP/TCGAbiolinksGUI
Operating system(s): Platform independent
Programming language: R
Other requirements: R*≥* 3.3
License: GNU GPL V3

## List of abbreviations

DEA: Differential expression analysis
DMR: Differentially methylated regions
ELMER: Enhancer Linking by Methylation/Expression Relationship
GDC: Genomic Data Commons
GUI: Graphical User Interface
LUSC: Lung Squamous Cell Carcinoma
MAE: Multi Assay Experiment
MAF: Mutation Annotation Format
TSS: Transcription start site

### Ethics approval and consent to participate

Not applicable

### Consent for publication

Not applicable

### Availability of data and material

Not applicable

### Author’s contributions

HN conceived the study. HN, BPB, MC and GB provided direction on the design of the Graphical User Interface. TCS developed and tested “GDC data”, “Epigenetic analysis”, “Clinical analysis”, “Genomic Analysis” and “Integrative analysis” interface. AC developed and tested the “Differential expression analysis” interface. CO developed and tested the “Network inference’ interface. TCS, AC and CO wrote the Users manual. TCS prepared the first draft of the manuscript. All authors were involved in the revision of the draft manuscript and have agreed to the final content.

## Acknowledgments

We are grateful to the OMICs lab and the GDC team for suggestions in the design of TCGAbiolinksGUI interface. We are also grateful for Susan MacPhee for critical review of the manuscript and vignettes.

## References

1. Gentleman, R.C., Carey, V.J., Bates, D.M., Bolstad, B., Dettling, M., Dudoit, S., Ellis, B., Gautier, L., Ge, Y., Gentry, J., et al.: Bioconductor: open software development for computational biology and bioinformatics. Genome biology 5(10), 80 (2004)

2. Colaprico, A., Silva, T.C., Olsen, C., Garofano, L., Cava, C., Garolini, D., Sabedot, T.S., Malta, T.M., Pagnotta, S.M., Castiglioni, I., Ceccarelli, M., Bontempi, G., Noushmehr, H.: Tcgabiolinks: an r/bioconductor package for integrative analysis of tcga data. Nucleic Acids Research 44(8), 71 (2016). doi:10.1093/nar/gkv1507. http://nar.oxfordjournals.org/content/44/8/e71.full.pdf+html

3. Chang, W., Cheng, J., Allaire, J., Xie, Y., McPherson, J.: Shiny: Web Application Framework for R. (2016). R package version 0.14. https://CRAN.R-project.org/package=shiny

4. Attali, D.: Shinyjs: Easily Improve the User Experience of Your Shiny Apps in Seconds. (2017). R package version 0.9.1. https://CRAN.R-project.org/package=shinyjs

5. Chang, W., Borges Ribeiro, B.: Shinydashboard: Create Dashboards with 'Shiny’. (2017). R package version 0.6.1. https://CRAN.R-project.org/package=shinydashboard

6. Pedersen, T.L.: shinyFiles: A Server-Side File System Viewer for Shiny. (2016). R package version 0.6.2. https://CRAN.R-project.org/package=shinyFiles

7. Yao, L., Shen, H., Laird, P., Farnham, P., Berman, B.: Inferring regulatory element landscapes and transcription factor networks from cancer methylomes. Genome biology 16(1), 105–105 (2015)

8. Chedraoui Silva, T., Coetzee, S.G., Yao, L., Hazelett, D.J., Noushmehr, H., Berman, B.P.: Enhancer linking by methylation/expression relationships with the r package elmer version 2. bioRxiv (2017). doi:10.1101/148726. http://www.biorxiv.org/content/early/2017/06/11/148726.full.pdf

9. Gu, Z., Eils, R., Schlesner, M.: Complex heatmaps reveal patterns and correlations in multidimensional genomic data. Bioinformatics (2016). doi:10.1093/bioinformatics/btw313. http://bioinformatics.oxfordjournals.org/content/early/2016/05/20/bioinformatics.btw313.full.pdf+html

10. Luo, W., Brouwer, C.: Pathview: an r/bioconductor package for pathway-based data integration and visualization. Bioinformatics 29(14), 1830–1831 (2013)

11. Mayakonda, A., Koeffler, P.H.: Maftools: Efficient analysis, visualization and summarization of maf files from large-scale cohort based cancer studies. BioRxiv (2016). doi:10.1101/052662

12. Kassambara, A., Kosinski, M.: Survminer: Drawing Survival Curves Using 'ggplot2'. (2017). R package version 0.4.0. https://CRAN.R-project.org/package=survminer

13. Gao, J., Aksoy, B.A., Dogrusoz, U., Dresdner, G., Gross, B., Sumer, S.O., Sun, Y., Jacobsen, A., Sinha, R., Larsson, E., et al.: Integrative analysis of complex cancer genomics and clinical profiles using the cbioportal. Science signaling 6(269), 1 (2013)

14. Cerami, E., Gao, J., Dogrusoz, U., Gross, B.E., Sumer, S.O., Aksoy, B.A., Jacobsen, A., Byrne, C.J., Heuer, M.L., Larsson, E., et al.: The cBio cancer genomics portal: an open platform for exploring multidimensional cancer genomics data. AACR (2012)

15. Turaga, N., Freeberg, M., Baker, D., Chilton, J., null, n., Nekrutenko, A., Taylor, J.: A guide and best practices for r/bioconductor tool integration in galaxy [version 1; referees: 1 approved, 1 approved with reservations]. F1000Research 5(2757) (2016). doi:10.12688/f1000research.9821.1

16. Grossman, R.L., Heath, A.P., Ferretti, V., Varmus, H.E., Lowy, D.R., Kibbe, W.A., Staudt, L.M.: Toward a shared vision for cancer genomic data. New England Journal of Medicine 375(12), 1109–1112 (2016)

17. Morgan, M., Obenchain, V., Hester, J., Pagès, H.: SummarizedExperiment: SummarizedExperiment Container. (2017). R package version 1.7.5

18. Sig, M.: MultiAssayExperiment: Software for the Integration of Multi-omics Experiments in Bioconductor. (2017). R package version 1.3.20. https://github.com/waldronlab/MultiAssayExperiment/wiki/MultiAssayExperiment-API

